# Subthalamic Nucleus dynamically encodes reward value and cognitive flexibility

**DOI:** 10.64898/2026.10.25.761735

**Authors:** Jennifer Collins, Grace Z. Zhou, Myles Reading, Jennifer L. Cornish, Asheeta A. Prasad

## Abstract

The subthalamic nucleus (STN) is classically associated with motor control but is increasingly recognised as a key component of the neural circuits underlying cognition. However, how the STN encodes learned reward associations and updates these representations when environmental contingencies change remains unclear. Here, we combined fibre photometry calcium imaging and chemogenetic inhibition to identify the contribution of the STN in reward learning and cognitive flexibility. During established Pavlovian conditioning, calcium imaging of the STN showed distinctive responses to reward predictive cue and reward delivery, while no response was detected to unrewarded cues. Upon reversal of reward predictive cues, STN activity initially remained associated with the previously rewarded cue before remapping to the newly rewarded cue as animals acquired the updated contingency. Chemogenetic inhibition of the STN impaired cue discrimination on established Pavlovian conditioning, delayed reversal learning, and disrupted outcome-specific Pavlovian-to-instrumental transfer. Together these results demonstrate that STN activity is required for flexible updating of learned reward associations and goal-directed behaviour.

These findings establish the STN as a dynamic integrative hub that encodes and updates learned reward representations to support cognitive flexibility, extending its role beyond motor control to include goal-directed behaviour. We show STN activity tracks reward-predictive cues and updates when contingencies reverse. Overall, these findings demonstrate that the STN is not just downstream of cortical control but actively supports updating of reward associations.

## Introduction

Adaptive behaviour requires the ability to form associations between environmental cues and outcomes, as well as the flexibility to modify these associations when contingencies change^1,2^. Although previously learned associations allow efficient responses to familiar situations, they can become maladaptive when environmental demands change. For example, when driving in a country with different road rules or driving conventions, individuals must draw on their existing driving knowledge while suppressing habitual response that are no longer appropriate and adapting to new rules. Such ability to adjust behaviour while overcoming interference from previously learned association, is a key aspect of cognitive flexibility^1–4^.

The subthalamic nucleus (STN) has classically been characterised as a key component of the basal ganglia motor circuit, associated by motor control and action selection^5–7^. More recently, however, the STN has emerged as an important regulator of cognitive control^8–11^. Anatomically, the STN is well positioned to integrate motor, associative, and limbic information. Indeed, fundamental studies have implicated that STN is involved in response inhibition, reward-related processing and regulation of cognitive control^12–14^.

For instance, electrophysiological studies have shown that distinct population of STN neurons encode appetite and aversive outcomes, as well as cues predicting reward, suggesting a role in representing reward-related information^15–18^. Lesion studies demonstrated that disruption of STN reduced impulsive choices and impaired Pavlovian conditioned responding^19–21^. In addition, pharmacological manipulation within the STN or its pathway have been shown to altered reward processing, drug-seeking reinstatement, and reacquisition of reward-seeking after extinction^22–24^.

Despite these growing evidence suggesting involvement of the STN in reward-related updating, how STN activity changes as animals acquire and update cue-reward associations remain less understood. It is also unclear whether STN activity represents reward-predictive cues or action-outcome association, and how these activities change when reward contingencies are reversed, and whether STN activity is necessary for these processes. Understanding these processes are critical, as flexible updating of reward related cues is fundamental for adaptive behaviour and is disrupted in several neuropsychiatric disorders. Indeed, the STN is a therapeutic target for deep brain stimulation for Parkinsons disease (PD), addiction, and treatment resistant obsessive compulsive disorder^25^. Better understanding of STN functions will improve clinical outcomes.

To address these questions, we combined fibre photometry and chemogenetics to investigate the contribution of the STN to reward learning, behavioural flexibility and motor behaviour across 5 experiments. Experiments 1 and 2 examined STN neural activity and its causal contribution during Pavlovian conditioning and Pavlovian Reversal to determine how reward-predictive cues and reward outcomes are encoded and updated. Experiment 3 and 4 investigated STN activity during instrumental learning and instrumental reversal, to determine how the STN encodes action-outcome association, and how these representations change following reversal, and whether STN activity is necessary for these action-outcome updating. Experiment 5 examined the effect of STN inhibition during reward probability using Pavlovian-to-instrumental Transfer (PIT), to determine whether the STN contributes to the influence of reward-predictive cues on instrumental behaviour. Locomotor behaviour was also assessed to examine potential motor effect of STN inhibition. Together, these experiments aimed to provide a comprehensive understanding of how the STN contributes beyond its traditional motor function, especially in representing and updating learned reward associations to support different form of behavioural flexibility and reward guided behaviour.

## Methods and Materials

### Subjects

Adult male Sprague Dawley rats (300-350g) were obtained from either the Animal Resource centre (Perth, Australia) or Ozgene (Perth, Australia). Subjects were housed in minimum groups of 2 in a climate-controlled colony room maintained on a 12:12 hour light/dark cycle (lights scheduled to turn on at 5:45 a.m). All behavioural procedures were conducted in the light cycle. All procedures were approved by the Animal Care and Ethics Committee at the University of Sydney and conducted in accordance with the National Health and Medical Research Code for the care and use of animals for scientific purposes in Australia.

### Surgeries

Stereotaxic surgeries were performed as previously described^26^ to unilaterally infuse viral vectors and implant fibre optic cannula for calcium recording, and to bilateral infuse of viral vector for chemogenetic inhibition. Rats were anesthetised in an induction chamber with 5% inhalant isoflurane and maintained at 2-3.5% while oxygen flow rates were maintained at 0.6L/min. Rats received a subcutaneous injection of carprofen (5mg/kg, Wagg & Purr), duplocillin (0.15mL/kg, Procaine (150 mg/mL) and Benzathine (115 mg/mL) penicillin, MSD Animal Health), and cefazolin (25mg/kg, AFT Pharmaceuticals) before being placed in the stereotaxic frame (Kopf Instruments). Bupivacaine (0.1mL, 0.5%, Pfizer) was administered at the incision site for local anaesthesia.

Subjects received stereotaxic surgery using flat skull coordinates relative to bregma, rat brain atlas (Paxinos & Watson, 2013). The coordinates targeting the STN were: −3.6 mm AP, ±2.4 mm ML, −7.9 mm DV for viral injection, and −3.6 mm AP, ±2.4 mm ML, −7.4 mm DV for fibre optic cannula (400 µm, NA 0.57, ceramic ferrule 2.5mm outer diameter, Thorlabs). Viral vectors were infused (unilaterally for calcium recording, bilateral for chemogenetic) using a 23-gauge, cone-tipped, 5µl stainless steel injector (SGE Analytical Science) connected to a bilateral infusion pump (UMP3 with SYS4 Micro-controller, World Precision Instruments, Inc., Sarasota, Florida, United States). Viral vectors were infused at a rate of 50 nL/min, with a total volume of 250 nL per hemisphere. The injector was left in place for an additional 5 minutes following infusion to allow viral diffusion before withdrawal. For fibre photometry recording, cannulas were implanted following viral injection. Four shallow burr holes were drilled into the frontal and parietal bone, and four jewellers’ screws (#0-80 x 1/8” Fine Thread Machine Screw Slotted Pan Head Stainless Steel 18-8, ASMC) were inserted into the skull to provide anchoring support. The skullcap was then secured using dental cement (cold-curing acrylic denture repair material, AVSCP0504000, AVSCV05000, Vertex Dental, Soesterberg, The Netherlands). Once the dental cement had cured, the incision was closed with sutures (3.0 Braided Silk, C7 needle, 18” black sutures, X786B, Look).

### Viral vectors

All viral vectors were purchased from Addgene. For fibre photometry experiments AAV5-syn-jGCaMP8f-WPRE was a gift from GENIE Project (500nL, Addgene viral prep # 162376-AAV5; http://n2t.net/addgene:162379; RRID: Addgene_162376; 100 µL at titer ≥ 7×10¹² vg/mL) used to monitor calcium activity. Vectors (250nl) were unilaterally infused into the STN, and following diffusion, a fibre optic cannula (200 µm, ThorLabs). For chemogenetic experiments, pAAV-hSyn-hM4D(Gi)-mCherry was a gift from Bryan Roth (500nL, Addgene viral prep # 50475-AAV5; http://n2t.net/addgene:50475; RRID:Addgene_50475; 100 µL at titer ≥ 7×10¹² vg/mL) and pAAV-hSyn-mCherry was a gift from Karl Deisseroth (500nL, Addgene viral prep #114472-AAV5; http://n2t.net/addgene:114472; RRID:Addgene_114472; 100 µL at titer ≥ 7×10¹² vg/mL) and were used for inhibition and control groups, respectively.

## Behavioural apparatus

### Behavioural chambers

Pavlovian conditioning, instrumental responding and PIT experiments were conducted in standard behavioural chambers (length = 29.35cm x width = 24.84cm x height = 18.67cm; Med Associates Inc. St Albans, VT, US, ENV-008-CT (Figure 1E). These chambers contained a recessed magazine port (4×4cm) that released sucrose pellets (45mg) (Able Scientific, ASF0023) from an attached pellet dispenser (Med Associates Inc. St Albans, VT, US, ENV-203M-45). A computer running MedPC-IV software controlled all events.

**Figure 1.**
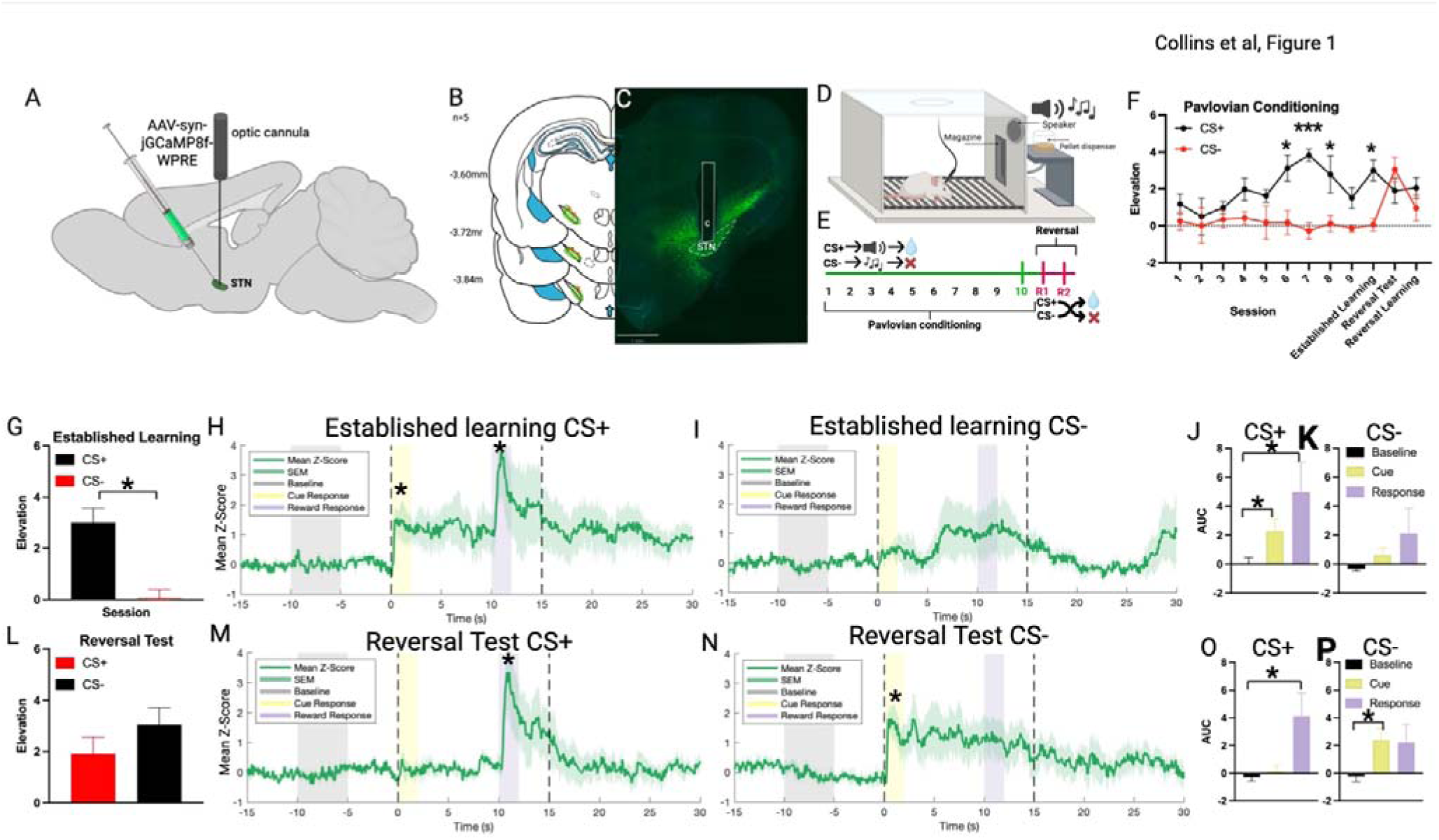
STN calcium activity during Pavlovian conditioning and Pavlovian Reversal. **(A)** Schematic of injection of AAV5-syn-jGCaMP8f-WPRE and cannula insertion into the STN. (B) Placement of cannula and viral expression for included subjects, n = 5. (C) Representative immunohistochemistry. (D) Schematic of Pavlovian conditioning and reversal paradigm. (E) Behavioural timeline, fibre photometry recordings occurred on day 10 Pavlovian training, reversal day 1 and 2 as indicated by the vertical line. (F) Magazine elevation across Pavlovian conditioning and reversal sessions for the rewarded cue (CS+) and non-rewarded cue (CS−). **G)** Magazine elevation during established Pavlovian conditioning on Day 10, showing responding to the CS+ and CS−. **(H–I)** STN calcium activity during presentation of the CS+ and CS−, respectively, during established Pavlovian conditioning (Day10). Mean *z*-scored calcium activity is shown by the green trace, with shading representing SEM. The baseline period is indicated in grey, cue onset at 0 s in yellow, and reward delivery at 10 s in purple. **(J–K)** Area under the curve (AUC) for baseline, cue-response, and reward-response periods during CS+ and CS− trials, respectively. **(L)** Magazine elevation during the initial reversal test following reversal of the CS–reward contingencies. **(M–N)** STN calcium activity during presentation of the new CS+ and new CS−, respectively, during the reversal test, shown in the same format as panels H–I. **(O–P)** AUC for baseline, cue-response, and reward-response periods during new CS+ and new CS− trials, respectively. All data are presented as mean ± SEM. Asterisks indicate significant differences relative to neural baseline (*p* < .05, *** *p* < .001).

### Open field locomotor chambers

Locomotor activity was assessed in a Noldus Phenotyper (length = 43.2 cm, width = 43.2 cm, height = 30.5 cm). Ethovision XT software (Noldus) was used to track and analyse distance travelled and average speed.

## Procedures

### Pavlovian Conditioning and Pavlovian reversal: Experiment 1 and 2

Experiments 1 and 2 aimed to investigate the contribution of the STN to reward learning and reversal of reward associated cues. Experiment 1 used fibre photometry to record STN activity during Pavlovian conditioning and Pavlovian reversal, whereas Experiment 2 used chemogenetic inhibition to test the causal contribution of the STN to these behaviours.

#### Pavlovian conditioning

Rats underwent 10 days of Pavlovian training, with 1 session each day. Each Pavlovian conditioning session consisted of 16 auditory CS-US presentations, including eight CS+ trials and eight CS− trials. The CSs were two distinct auditory cues, a tone (1500 Hz) and a clicker (90 Hz) that last 15 second. CS identity was counterbalanced across rats, such that either the tone or clicker served as the CS+, other served as CS-.

During rewarded trials, the CS+ was paired with delivery of a sucrose pellet at the 10 second; where CS- has no consequence. The order of CS presentations varied across sessions, and inter-trial intervals were randomly selected, with an average duration of 4 min. Each session lasted 70 mins. CS identity and magazine entries were recorded using MED-PC IV software.

To assess STN activity and its causal contribution to established Pavlovian reward learning, fibre photometry recordings and chemogenetic inhibition were conducted on Day 10 of Pavlovian conditioning. This allowed STN activity and function to be examined once animals had acquired stable discrimination between the rewarded and non-rewarded cues.

#### Pavlovian Reversal

Following Pavlovian conditioning, rats underwent Pavlovian reversal. During reversal, the cue-outcome contingencies were switched, such that the previous CS+ became the CS−, and the previous CS− became the CS+. In experiment 1, fibre photometry calcium recordings were conducted on the first and second day of reversal, following 10 days of Pavlovian training (Figure 1E) In experiment 2, rats received an additional three days of Pavlovian training without chemogenetic manipulation before reversal training commenced. This was to minimise any potential residual manipulation effects from Day 10 Pavlovian training. Chemogenetic inhibition was then conducted on the first day of the Pavlovian Reversal training (Figure 2B).

**Figure 2.**
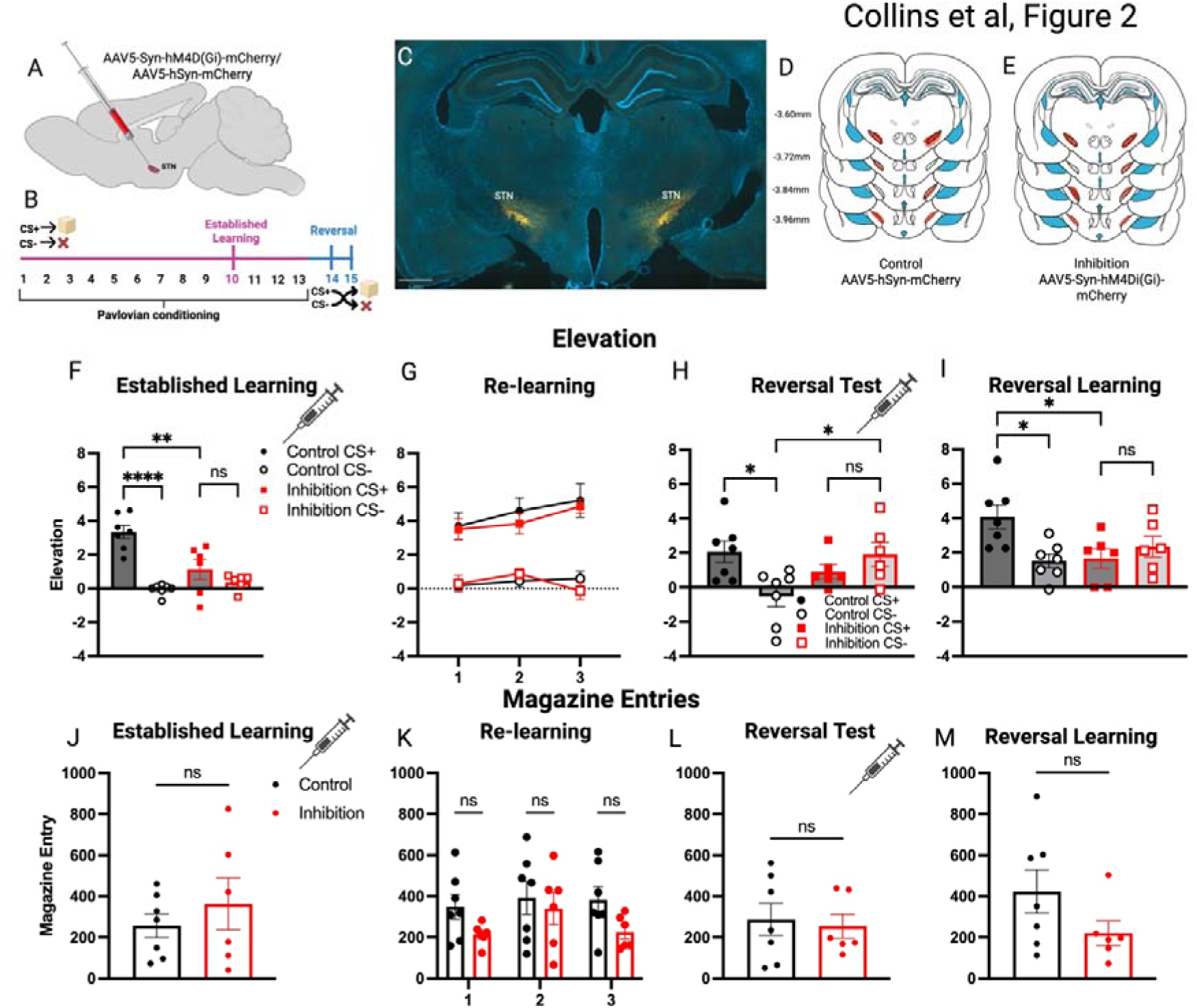
Chemogenetic inhibition of the STN during Pavlovian conditioning and reversal. **(A)** Schematic showing placement AAV5-hSyn-mCherry/-hM4D(Gi)-mCherry. **(B)** Experimental timeline for Pavlovian conditioning, established learning, retraining, and reversal testing. Chemogenetic inhibition was administered during the established-learning and initial reversal-test sessions, as indicated by the syringe symbols. **(C)** Representative image of viral in the STN. **(D-E)** Viral expression and injection-site distribution for the control and inhibition groups, respectively. **(F)** Magazine elevation during established Pavlovian learning (day 10) for the control and inhibition group. **(G)** Magazine elevation across the 3 days of retraining preceding reversal. **(H)** Magazine elevation during the initial reversal test, in which CS+ and CS- contingency swapped. **(I)** Magazine elevation during the subsequent reversal-learning session conducted on the following day without chemogenetic manipulation (**J)** Total magazine entries during the established-learning test. **(K)** Total magazine entries across the 3 days of retraining. **(L)** Total magazine entries during the initial reversal test. **(M)** Total magazine entries during the subsequent reversal-learning session. Control and inhibition groups are shown in black and red, respectively. All data are presented as Mean ±SEM. Significant effect is indicated by (* p <0.05; ** p<0.01; **** p<0.00001; ns, not significant).

### Instrumental conditioning and Reversal: Experiment 3 and 4

Experiments 3 aimed to investigate the contribution of the STN in action outcome reward learning and flexible updating. Experiment 3 used calcium fibre photometry to record STN activity during instrumental conditioning and instrumental reversal. Followed by chemogenetic inhibition of the STN during instrumental conditioning in experiment 4.

#### Instrumental Conditioning

Rats first underwent four x 20-minute sessions of magazine training over two days. During each session, 10 sucrose pellets were randomly delivered to the magazine with no lever presentation. This was to acclimatise animals to behavioural chambers and to seek the reward in the magazine ^26^. Instrumental conditioning then commenced with both rewarded and non-rewarded levers available. Each rewarded lever resulted in one sucrose-pellet delivery (Fixed Ratio one (FR1)), followed by a 10-second timeout during which additional responses were not reinforced. Presses on the rewarded lever within the time out were not reinforced. Press on the non-rewarded lever had no programmed consequences. Rewarded and non-rewarded lever were assigned consistently within each rat and counterbalanced between rats. All rats completed at least ten x 30-minute instrumental-training sessions before neural recording or manipulation occurred. Learning was assessed by discrimination between the rewarded and non-rewarded lever presses, with greater responding on the rewarded lever than non-rewarded lever indicating successful acquisition. Fiber photometry recording and chemogenetic inhibition was then conducted at the Day 10 of the instrumental training.

#### Instrumental reversal

Following 10 days of instrumental conditioning, rats underwent reversal learning in which the rewarded and non-rewarded lever contingencies were reversed for each rat. Reversal training continued for 10 days until rats demonstrated the updated contingency by preferentially responding on the new rewarded lever. For fibre photometry (Experiment 3), calcium recording was recorded on the 1^st^ day of the reversal (reversal test), 2^nd^ day of the reversal (reversal learning) and 10^th^ day of the reversal (established reversal) (Figure 3F). This paradigm assessed cognitive flexibility by measuring the ability to adapt behaviour following changes in reward contingencies^27^.

**Figure 3.**
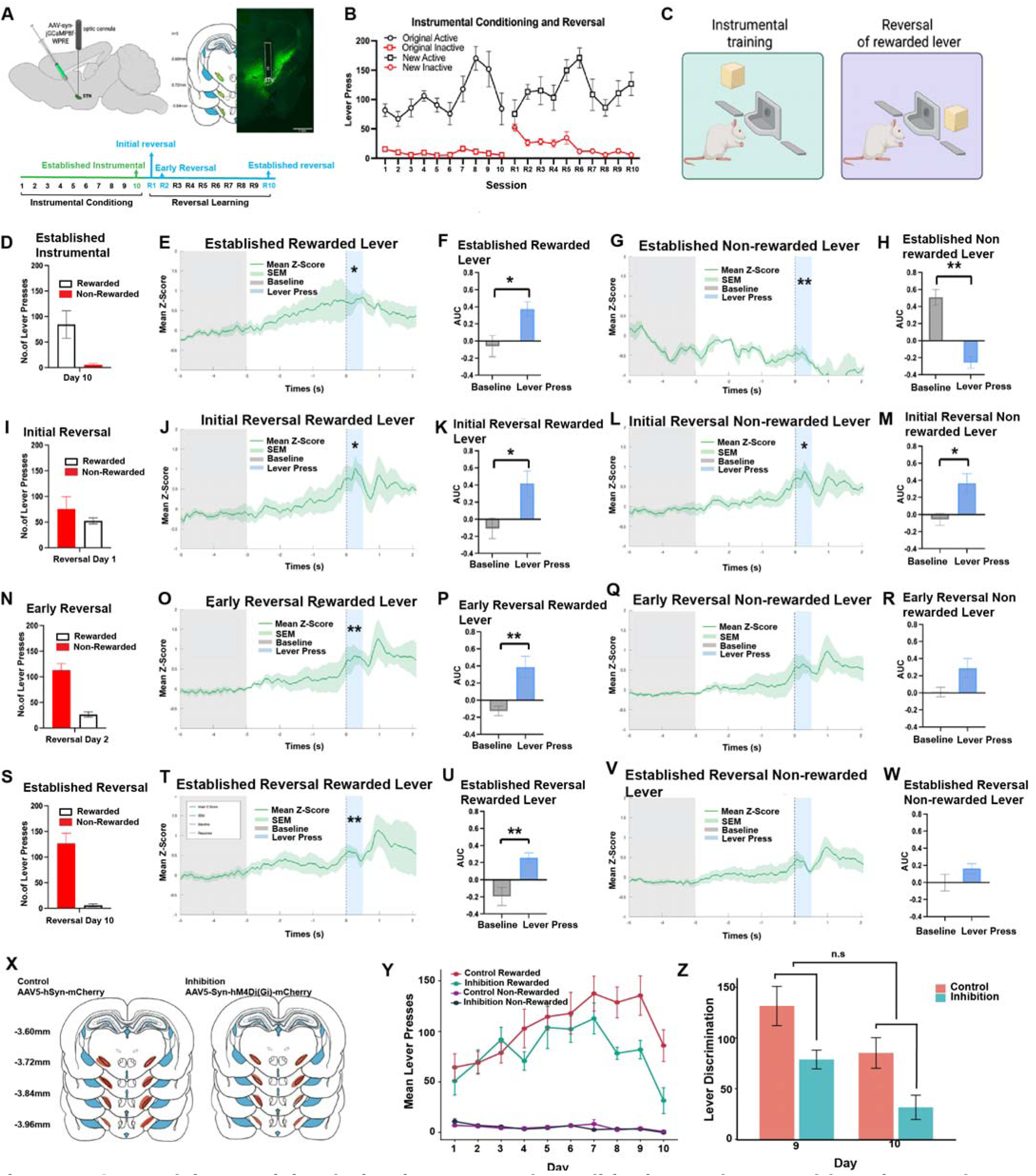
STN calcium activity during instrumental conditioning and reversal learning, and effects of chemogenetic inhibition on instrumental responding. **(A)** Experimental design for fibre photometry recordings. Left, schematic showing injection of AAV-syn-jGCaMP8f-WPRE and implantation of an optic fibre targeting the STN. Middle, representative viral expression and optic-fibre placement in the STN. Right, locations of viral expression and optic-fibre placements for included rats (*n* = 5). Bottom, experimental timeline indicating fibre photometry recording sessions during established instrumental conditioning (Day 10), initial reversal (Reversal Day 1), early reversal learning (Reversal Day 2), and established reversal learning (Reversal Day 10). **(B)** Lever-press responses across instrumental conditioning and reversal learning. During instrumental conditioning, responses on the original rewarded and non-rewarded levers are shown. Following reversal of the lever–outcome contingency, responses on the newly rewarded and non-rewarded levers are shown. **(C)** Schematic illustrating the behavioural contingencies during instrumental training and following reversal of the rewarded lever. **(D–H) Established instrumental conditioning. (D)** Rewarded and non-rewarded lever presses during established instrumental conditioning. **(E)** Mean *z*-scored STN calcium activity aligned to rewarded-lever presses. **(F)** Area under the curve (AUC) during the baseline and lever-press periods for rewarded-lever presses. **(G)** Mean *z*-scored STN calcium activity aligned to non-rewarded-lever presses. **(H)** AUC during the baseline and lever-press periods for non-rewarded-lever presses. **(I–M) Initial reversal. (I)** Rewarded and non-rewarded lever presses during the initial reversal session. **(J)** Mean *z*-scored STN calcium activity aligned to rewarded-lever presses. **(K)** AUC during the baseline and lever-press periods for rewarded-lever presses. **(L)** Mean *z*-scored STN calcium activity aligned to non-rewarded-lever presses. **(M)** AUC during the baseline and lever-press periods for non-rewarded-lever presses. **(N–R) Early reversal learning. (N)** Rewarded and non-rewarded lever presses during early reversal learning. **(O)** Mean *z*-scored STN calcium activity aligned to rewarded-lever presses. **(P)** AUC during the baseline and lever-press periods for rewarded-lever presses. **(Q)** Mean *z*-scored STN calcium activity aligned to non-rewarded-lever presses. **(R)** AUC during the baseline and lever-press periods for non-rewarded-lever presses. **(S–W) Established reversal learning. (S)** Rewarded and non-rewarded lever presses during established reversal learning. **(T)** Mean *z*-scored STN calcium activity aligned to rewarded-lever presses. **(U)** AUC during the baseline and lever-press periods for rewarded-lever presses. **(V)** Mean *z*-scored STN calcium activity aligned to non-rewarded-lever presses. **(W)** AUC during the baseline and lever-press periods for non-rewarded-lever presses. **(X)** Viral-expression locations for the control and chemogenetic inhibition groups. **(Y)** Mean rewarded- and non-rewarded-lever presses across instrumental-conditioning days in the control and inhibition groups. **(Z)** Lever-discrimination scores, calculated as rewarded minus non-rewarded lever presses, on Days 9 and 10 for the control and inhibition groups. The Day × Group interaction was not significant, indicating that the change in lever discrimination from Day 9 to Day 10 did not significantly differ between groups. Data are presented as mean ± SEM. Shaded regions represent SEM. *p* < .05, \**p* < .01; n.s., not significant.

### Pavlovian-Instrumental Transfer (PIT): Experiment 5

Chemogenetic manipulation was applied to PIT paradigm to investigate whether STN is sensitive to reward probability, and how these uncertain predictive cues influence goal-direct behaviour. The PIT procedure consisted of three stages: Pavlovian conditioning, instrumental training and the PIT test.

#### Pavlovian conditioning: Reward probability under PIT

Rats first underwent 10 days of Pavlovian conditioning, during which two auditory conditioned stimuli (CSs) were associated with different probabilities of sucrose pellet delivery. One cue was paired with reward delivery on 70% of trials (CS70), whereas the other cue was paired with reward delivery on 30% of trials (CS30). Each Pavlovian conditioning session lasted approximately 1 hour, and each cue was presented 10 times per session. Cue presentation order was counterbalanced across rats and sessions (Figure 4D).

**Figure 4.**
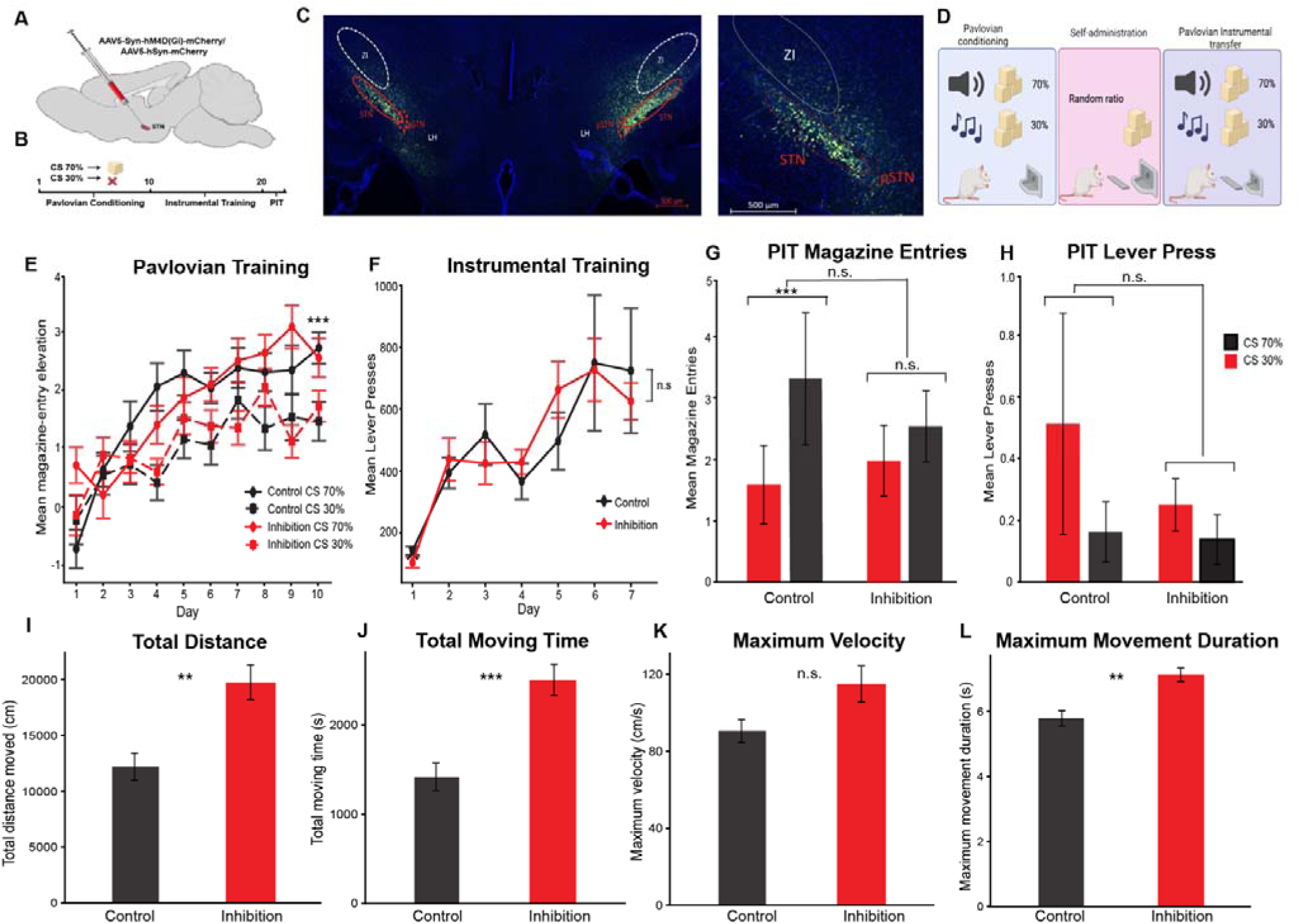
Chemogenetic inhibition of the STN during Pavlovian to instrumental transfer. **(A)** Schematic showing injection of AAV5-hSyn-hM4D(Gi)-mCherry or AAV5-hSyn-mCherry into the STN. **(B)** Experimental timeline showing Pavlovian conditioning, instrumental training, and the Pavlovian-to-instrumental transfer (PIT) test. **(C)** histological images showing viral expression in the STN. **(D)** Schematic of the behavioural paradigm. During Pavlovian conditioning, rats received a 70% CS and a 30% CS associated with different reward probabilities, followed by instrumental self-administration training and the subsequent PIT test. **(E)** Mean magazine-entry elevation across 10 days of Pavlovian conditioning for the 70% CS and 30% CS in control and inhibition groups. Rats showed greater magazine-entry elevation to the 70% CS than to the 30% CS, with no significant group difference or CS type × Group interaction. **(F)** Mean lever presses across instrumental training. Lever pressing increased across 7 days of training, indicating acquisition of the instrumental response, with no significant difference between the control and inhibition groups. **(G)** Mean magazine entries during presentation of the 70% CS and 30% CS in the PIT test. Control rats showed significantly greater magazine entries during presentation of the 70% CS than the 30% CS, whereas this difference was not significant in inhibition rats. **(H)** Mean lever presses during presentation of the 70% CS and 30% CS in the PIT test. Lever pressing did not differ significantly by CS type, group, or the CS type × Group interaction. **(I–L)** Locomotor activity in control and inhibition rats, quantified as total distance moved **(I)**, total moving time **(J)**, maximum velocity **(K)**, and maximum movement duration **(L)**. All data are presented as mean ± SEM. Significance is indicated by **p < .01, ***p < .001; ns, not significant.

#### Instrumental conditioning

Following Pavlovian conditioning, rats underwent instrumental training, during which they learned to press a lever to obtain sucrose pellets. Two levers were available, but only one lever resulted in reward delivery. The rewarded lever was counterbalanced across rats. Rats received two x 30-minute instrumental training sessions per day across 7 days. On Day 1, responding was reinforced on a fixed ratio 1 schedule, such that each rewarded lever press resulted in one sucrose pellet delivery. From Days 2–4, rats were trained on a random ratio 5 schedule, in which rewarded lever presses had a pseudorandomised 20% chance of producing reward. From Days 5–7, the reinforcement schedule was shifted to a random ratio 10 schedule, in which rewarded lever presses had a pseudorandomised 10% chance of producing reward (Figure 4D).

#### PIT test

Before the PIT test, rats received one instrumental extinction and one Pavlovian reminder session. Conditioned cues were presented in the same manner as the Pavlovian conditioning, while both levers were available. No rewards were delivered during the testing session. Chemogenetic inhibition was conducted during the PIT test to assess its effect on cue-driven instrumental responding.

#### Fibre Photometry (Experiment 1 and 3)

To conduct neural recordings, subjects were tethered to the fibre photometry rig via a subject cable (Ø400µm, 0.57NA, Doric Lenses, Canada) fitted to their cannula with a mating sleeve (ADAF1-5, ThorLabs, USA). A real-time signal processor (RZ10X Lux-I/0 Processor, Tucker-Davis Technologies, USA) transmitted light at 465nm and 405nm to excite jGCaMP8f. The 465nm LED stimulated calcium dependent fluorescent signals, while the 405nm LED was utilized for calcium fluorescence, providing a negative internal isosbestic control ^28^.

The LED driver sinusoidally modulated LEDs at 209Hz for the 465nm LED and 331Hz for the 405nm LED, maintaining light intensity between 10-30 µW to minimise photo-bleaching. Driver cables (Ø200µm, 0.5 NA, LXFX-KIT-2C, Tucker-Davis Technologies, USA) sent light through the Doric Mini Cube (FMC6_IE, Doric Lenses) to the subject. Fluorescence activity was collected and transmitted to the RZ10x photosensor via response cables (Ø600µm, 0.5 NA, LXFX-KIT-2C, Tucker-Davis Technologies, USA). The photosensor low-pass then filtered the data at 6Hz to remove noise and demodulated 405nm and 465nm signals. Synapse software (Tucker-Davis Technologies, USA) controlled the RZ10x and recorded signals. Behavioural events (cue-onset) were time-locked to fibre photometry data by Synapse for data analysis.

#### Chemogenetic manipulation (Experiment 2, 4 & 5)

All chemogenetic manipulations conducted at least 6 weeks following surgery. Rats received an intraperitoneal injection of clozapine (2.5 mg/kg; #C6305, Sigma Aldrich); administered at a volume of 1 mL/kg. Clozapine was diluted with 5% DMSO and saline and delivered 30min prior to behavioural testing.

#### Immunohistochemistry

Following the conclusion of behavioural experiments, rats were deeply anesthetised with Lethabarb (100mg/kg, i.p., Virdac). Rats were transcardially perfused with 150 ml of 0.9% saline, then followed by 500ml of 4% paraformaldehyde (PFA), in 0.1 M phosphate buffer (PB), pH 7.4. Brains were extracted and postfixed in 4% PFA for at least 2 hours in the same fixative and then stored in a 20% sucrose solution with phosphate-buffered saline (PBS) overnight for sufficient cryoprotection. Brains were frozen and then coronally sectioned at 40 µm using cryostat (NX50, Cryostar Expredia). Four serially adjacent sets of tissue were collected and stored in anti-freeze solution (30% glycerol, 20% ethylene glycol in 0.1M phosphate buffer, pH 7.4). Tissue sections were washed with 0.1M tris-buffered saline (TBS) and blocked with casein/bovine serum blocking buffer (1% casein, 4% bovine serum albumin, 1.5% glycine, 0.05% Triton X100, 0.02% sodium azide in 0.1M PBS) for one-hour to prevent non-specific antibody binding. Sections were incubated overnight with primary rabbit anti-mCherry to identify viral expression for calcium signalling experiments, and chemogenetic experiments respectively (1:500, mCherry polyclonal antibody, PA5-34974, Invitrogen). After washing with TBS with 0.05% Tween, sections were incubated with Alexa Fluor 488-conjugated secondary antibody (1:500, Alexa Fluor 488 goat anti-rabbit (A-11034), ThermoFisher Scientific) for 3-hours. Sections were counterstained with Hoechst (1:500, Hoechst 33342, 62249, ThermoFisher Scientific) to visualise cell nuclei. Sections were mounted onto Superfrost glass slides (EPBRSF41296, Expredia) and cover-slipped with mounting media (Prolong Diamond Antifade Mountant, P36970, Invitrogen). Slide scanner VS200-MTL (Olympus) was then used to visualise viral expression and respective cannula placements within the STN.

### Data processing and analyses

#### Behavioural Data

For Pavlovian Conditioning, cue-evoked magazine entry was quantified as an elevation score. This was calculated as the number of magazine entries during the 15 s CS period minus the number of magazine entries during the 15 s pre-CS baseline period. With higher number indicate stronger conditioned toward the cue. For instrumental responses the number of lever presses and magazines entries are recorded. For PIT, behavioural responding was assessed using the number of lever presses and magazine entries during the 15 s CS period. No elevation of score is used as PIT is essentially an extinction, with baseline level decrease throughout the test. Higher levels of CS- evoked lever pressing are interpreted as stronger transfer.

Statistical analyses were conducted using SPSS Statistics 28 (IBM, NY, USA), RStudio (version 2026.01.1+403). Assumptions for ANOVA and linear mixed models were assessed prior to analysis. Where the assumption of sphericity was violated, Greenhouse–Geisser corrections were applied. Graphs were generated using Python and MATLAB version 24.2.0.2773142 (R2024b; The MathWorks Inc., Natick, MA, USA), and Prism 9 (GraphPad Software).

#### Fibre photometry Data

Analyses of fibre photometry data was completed using Custom MATLAB scripts. Time ranges in the raw data stream with motion artifacts were manually excluded. High-frequency noise was removed through low-pass digital filtering at 2 Hz. To control for calcium fluorescence and to normalise data, ΔF/F (change in fluorescence relative to baseline) was calculated ^28^. The control 405nm signal was linearly fit to the 465nm signal. ΔF/F was calculated using the formula (465nm signal - fitted 405nm signal) / (fitted 405nm signal), yielding the final signal for further statistical analysis.

For Pavlovian conditioning sessions calcium signals were averaged around behavioural events: CS+ (average of all CS+ events in one session) and CS- (average of all CS- events in one session) and compared to the neural baseline. Z-scores were calculated on a trial-by-trial basis using the formula (all data points ΔF/F- baseline mean ΔF/F) / (baseline standard deviation ΔF/F) where baseline was between 10 and 5 seconds pre cue onset and all data points were during the 15 second auditory cue onset. This period includes the 10 seconds of auditory cue and on the 10^th^ second the pellet is delivered in the case of CS+. The area under the curve (AUC) of the calcium signal was analysed per subject surrounding CS presentation and reward delivery, then averaged across all subjects.

Group mean of the Z-score signal surrounding behavioural events was determined. Group mean of the Z-score signal surrounding behavioural events was determined for each session. To examine behaviour-related increases in activity AUC, the group mean Z-scores were computed with trapezoidal numerical integration. For Pavlovian task, the response to auditory cue and reward was detailed with the baseline of −10 to 5 seconds prior to cue onset, 0 to 2 seconds, and 10 to 12 seconds for auditory cue response and reward consumption, respectively.

To assess calcium activity during recorded sessions, two-way repeated-measures ANOVAs were performed on mean Z-score AUCs across four epoch (baseline, cue response, reward response) and two CS type (CS+ vs CS−). Post-hoc analyses were performed using Tukey’s tests where required.

For instrumental conditioning sessions, epochs were created around lever presses on the rewarded and non-rewarded levers. Lever-press events were temporally isolated such that the photometry signal associated with one examined event did not overlap with that of another. For each event, photometry data was extracted from −10 to +10 s relative to the lever press. Z-scores were calculated on a trial-by-trial basis as (ΔF/F − baseline mean ΔF/F) / baseline standard deviation ΔF/F, using the −10 to −5 s period before each lever press as baseline. To assess STN calcium activity during instrumental conditioning and reversal learning, the area under the curve (AUC) of the calcium signal surrounding rewarded and non-rewarded lever presses was calculated. Paired *t*-tests were used to compare calcium activity during the lever-press response period with the corresponding baseline period.

## Results

### Experiment 1: Calcium fibre photometry recording in the STN during Pavlovian condition and Pavlovian reversal

#### Histological validation

To assess viral expression of jGCaMP8f and correct placement of the optic fibre cannula, brains sections were histologically processed and analysed with reference to Paxinos and Watson (2006) description of STN coordinates. One subject was excluded from this experimental cohort due to incorrect cannula placement, resulting in n=5 for analysis (Figure 1A-C).

#### Pavlovian conditioning and reversal behavioural response and STN Calcium activity

Behaviourally, elevation scores changed significantly across the 10 days of Pavlovian training, *F*(11, 88) = 2.185, *p* = 0.020. By day 10, elevation scores were significantly higher during the CS+ than CS−, *t*(4) = 7.59, *p* = 0.001 (Figure 1D-F), indicating that rats had learned to discriminate between rewarded and non-reward cues. During the fibre-photometry recording session on day 10, STN calcium activity significantly increased from the pre-cue baseline for both the cue-evoked (auditory CS+ onset), (*t*(8) = 2.448, *p* = 0.04) and reward-evoked period (sucrose pellet delivery) (*t*(8) = 2.315, *p* = 0.04). Whereas no significant difference was observed from baseline to auditory CS- onset, cue-evoked period (*t*(8) = 2.053, *p* = 0.070) and to reward-evoked period (*t*(8) = 1.424, *p* = 0.192) (Figure 1G-K). These findings indicate that STN activity was selectively associated with the presentation of an established reward-predictive cue and the subsequent reward outcome, but not with the presentation of an unrewarded cue.

During Pavlovian reversal, the previously established CS– outcome contingencies were reversed. Elevation scores did not significantly differ between the new CS+ (previously the CS−) and the new CS− (previously the CS+) on either the first day (*t*(4) = 0.97, *p* = 0.390), or the second day (*t*(4) = 0.97, *p* > 0.390). Suggesting that the rats had not yet behaviourally discriminated between the reversed cue-outcome contingencies during early reversal learning.

The STN calcium activity during the new CS+ presentation on the first day of the reversal does not differ significantly from baseline period, *t*(8) = 0.74, *p* = 0.482, but a higher significant calcium activity response was observed during reward delivery period (compared to baseline period), *t*(8) = 2.62, *p* = 0.031. In contrast, the new CS−, which had previously served as the rewarded CS+, produced a significant cue-evoked response relative to baseline, *t*(8) = 2.74, *p* = .025, but not when the sucrose pellet was delivered, *t*(8) = 1.77, *p* = .116. These findings suggest that during early reversal learning, the STN continued to respond to the cue that had previously predicted reward, despite the reversal of its contingency. Interestingly, STN activity responded to reward delivery following the new CS+ before a clear cue-evoked response to the new reward-predictive cue had developed (Figure 1L-P).

### Experiment 2: Chemogenetic inhibition of the STN during Pavlovian condition and reversal behaviour

#### Histological validation

Only subjects with correct bilateral placement were included in analyses. Three subjects were excluded from both the control and inhibition group due to incorrect viral expression, resulting in n=7 and n=6 respectively chemogenetics inhibition of STN during Pavlovian conditioning and reversal experiment analysis (Figure 2A-E).

#### Chemogenetic inhibition of Pavlovian and reversal behavioural

Across 9 days of Pavlovian conditioning, rats increasingly responded for CS+ than to the CS- across sessions, *F*(1, 10) = 102.1, *p* < .0001. On day 10, where chemogenetic manipulation was administered, rats in control groups showed a significantly higher elevation during the CS+ than during the CS- (*t*(6) = 7.66, *p* < .0003), indicating clear discrimination between the two cues. In contrast, rats that received chemogenetic inhibition showed no significant difference in elevation between CS+ and CS- (*t*(5) = 1.53, *p* = .186), (Figure 2F). To further understand whether this is impacted by a more general reduction in magazine-directed responding, we also analysed the number of magazine entries, where no significant difference was found between control and inhibition group (*t*(11) = 0.81, *p* = .44) (Figure 2J). Together, these findings suggesting STN inhibition impairs discrimination between rewarded and non-rewarded cue, without producing a general reduction in magazine-directed responding.

Before reversal testing, rats underwent three days of retraining with the original Pavlovian cue - reward contingencies to minimise any residual effects of the previous chemogenetic manipulation. By the end of retraining, no significant group differences were observed in elevation scores, *F*(1, 11) = 0.344, *p* = .57 (Figure 2G), or total magazine entries, *t*(11) = 0.33, *p* = .75 (Figure 2K), indicating comparable behavioural performance between the control and inhibition groups before reversal test.

Following retraining, rats underwent a reversal test under chemogenetic inhibition in which the CS+ and CS- contingencies were switched. Control rats showed significantly greater elevation scores compared to the new CS+ to the new CS−, *t*(6) = 2.97, *p* = .020, whereas rats that received chemogenetic inhibition of the STN showed no significant difference between the cues, *t*(5) = 1.22, *p* = 0.28. On the following day, rats completed a second reversal-learning session without chemogenetic manipulation. This allowed to assess whether the behavioural effect observed during STN inhibition persisted after the manipulation was no longer present. Control rats continued to show significant discrimination between the new CS+ and CS−, *t*(6) = 3.97, *p* = .007, whereas no significant cue discrimination was observed in the STN-inhibition group, *t*(5) = 0.62, *p* = .56 (Figure 2H). Again, no significant differences between groups were observed in total magazine entries during either the initial reversal test, *t*(11) = 0.33, *p* = .14 (Figure 2L), or the subsequent reversal-learning session, *t*(10) = 1.40, *p* = 0.190 (Figure 2I-M). Together, these findings suggest that STN inhibition impaired flexible updating of cue-reward associations without producing a general reduction in magazine-directed responding.

### Experiment 3: Calcium fibre photometry recording in the STN during instrumental training and reversal

#### Histological validation

Only subjects with correct jGCaMP8f expression and cannula placement were included for analysis, n=5. During instrumental training and reversal, lever-pressing behaviour and STN calcium activity were examined (Figure 3A). Analyses focused on four sessions: established instrumental performance (Day 10; the final day of instrumental training), initial reversal (Day 11; the first reversal session), early reversal learning (Day 12; the second reversal session), and established reversal performance (Day 20; the tenth reversal session), (Figure 3 D-F). Calcium activity was analysed around both lever presses and magazine entries. Lever presses were classified according to whether they occurred on the rewarded or non-rewarded lever, and behavioural events were further separated according to whether they were followed by reward delivery.

#### Established instrumental performance

By Day 10 of instrumental training, rats pressed the rewarded lever significantly more often than the non-rewarded lever, *t*(8) = 2.92, *p* = .020 (Figure 3D). For the rewarded lever presses that were followed by a reward delivery, calcium activity during the lever presses period was significantly higher from baseline, *t* (8) = 2.86, *p* = .020. (Figures 3E-F). For non-rewarded lever presses, calcium activity was significantly lower from baseline during the lever-press period, *t*(4) = 6.63, *p* = .003 (Figures 3 G-H).

#### Initial reversal

Following instrumental training, rats underwent reversal of rewarded and non-rewarded lever contingencies. Behaviourally, rats did not significantly differ in number of lever presses on the newly rewarded lever (previous non-rewarded lever) and the newly non-rewarded lever (previous rewarded lever) *t*(8) = 0.09, *p* = .38 (Figure 3L).

The calcium activity of lever presses on the newly rewarded lever and the previously rewarded lever were both significantly higher from baseline, *t*(8) = 2.78, *p* = .020; *t*(8) = 3.18, *p* = .010, respectively (Figures 3 J-M).

#### Early reversal learning

On the following day, during the second reversal session, rats pressed the newly rewarded lever significantly more often than the newly non-rewarded lever, *t*(8) = 6.39, *p* = .0002 (Figure 3N), demonstrating behaviour had shifted toward the lever associated with the new reward contingency.

Calcium activity during presses on the newly rewarded lever was significantly higher to baseline, *t*(8) = 3.70, *p* = .006 (Figure 3 O-P). In contrast, calcium activity during presses on the previously rewarded lever, which was now non-rewarded, did not significantly differ from baseline during the lever-pressing period *t*(8) = 2.21, *p* = .058 (Figures 3Q-R).

#### Established reversal performance

On the Day 10 of the reversal learning, rats pressed the newly rewarded lever significantly more often than the newly non-rewarded lever, *t*(8) = 6.08, *p* = .0003 (Figure 3S).

For presses on the reversed rewarded lever, calcium activity was significantly higher from baseline during the lever-pressing period, *t*(8) = 3.80, *p* = .006 (Figure 3T-U). In contrast, calcium activity during pressing the non-rewarded lever did not significantly differ from baseline period, *t*(8) = 1.47, *p* = .180 (Figure 3 V-W).

### Experiment 4: Chemogenetic inhibition of the STN during established instrumental conditioning

#### Histological validation

Only subjects with correct bilateral placement were included in analyses. Sixteen subjects were excluded from both the control and inhibition group due to incorrect viral expression, resulting in n=13 for control and n=11 for inhibition analysis for chemogenetics inhibition of STN during Instrumental conditioning. (Figure 3X)

#### Chemogenetic inhibition of Instrumental conditioning

Lever discrimination was quantified as the difference between rewarded and non-rewarded lever presses (rewarded − non-rewarded). Lever discrimination significantly increased from Day 1 to Day 9, *F*(1, 22) = 30.62, *p* < .001. This increase in lever discrimination across training did not significantly differ between the control and inhibition groups, *F*(1, 22) = 3.02, *p* = .10. (Figure 3Y)

On Day 10, rats underwent chemogenetic inhibition during instrumental responding. Following exclusion of animals with incorrect viral expression, lever discrimination appeared to decrease from Day 9 to Day 10. To determine whether this decrease was specifically associated with chemogenetic inhibition, discrimination scores from Days 9 and 10 were analysed using a mixed two-way ANOVA, with Day as a within-subject factor and Group as a between-subject factor.

Averaged across Days 9 and 10, lever discrimination significantly differed between the control and inhibition groups, *F*(1, 22) = 9.50, *p* = .005. Across both groups, lever discrimination was significantly lower on Day 10 than on Day 9, *F*(1, 22) = 14.09, *p* = .001. This reduction in lever discrimination from Day 9 to Day 10 did not differ between the control and inhibition groups *F*(1, 22) = 0.001, *p* = .975. Together, these findings provide no evidence that chemogenetic inhibition selectively altered lever discrimination (Figure 3Z).

### Experiment 5: Chemogenetic inhibition of the STN during PIT

#### Histological validation

Only rats with confirmed bilateral placement within the STN were included in analyses, resulting in n=8 per group. Chemogenetic inhibition was conducted on PIT testing day (Figure 4A-D).

#### Pavlovian Training

On day 10 of Pavlovian training, rats showed stronger conditioned responding to the cue associated with the higher reward probability, where magazine-entry elevation was significantly higher for the CS 70% than for the CS 30%, *F*(1, 14) = 26.11, *p* < .001. There was no significant difference between two groups, *F*(1, 14) = 0.07, *p* = .788). The difference in elevation between CS 70% and CS 30% did not differ significantly between groups, *F*(1, 14) = 0.15, *p* = .703 (Figure 4E).

#### Instrumental Training

Rats made significantly more rewarded lever presses on Day 7 than on Day 1, *F*(1, 14) = 29.57, *p* < .001. This suggests that rats acquired the instrumental response across instrumental training. The overall rewarded lever pressing did not significantly differ between the Control and Inhibition groups, *F*(1, 14) = 0.36, *p* = .558. The increase in rewarded lever pressing from Day 1 to Day 7 did not significantly differ between groups, *F*(1, 14) = 0.09, *p* = .769. (Figure 4F).

#### Pavlovian to Instrumental Transfer

For magazine entries during PIT, rats made more magazine entries during the high reward probability CS (CS 70%) than during the low reward probability CS (CS 30%) (*b* = 1.725, *p* < .001). No significant difference in magazine entries was found between the Control and Inhibition groups (*b* = 0.387, *p* = .699). The increases in magazine entries during the CS 70% was more pronounced in control rats than in Inhibition rats (*b* = −1.163, = .04). Follow-up simple-effects comparisons confirmed that control rats made significantly more magazine entries during the CS 70% than the CS 30%, *t*(302) = 4.25, *p* < .001. In contrast, the difference in magazine entries between CS70% and CS30% was not significant in the Inhibition group, *t*(302) = 1.39, *p* = .167 (Figure 4G)

For lever pressing during the PIT test, there was no significant difference in number of lever pressing during presentation of CS 70% compared with the presentation of CS 30% (*b* = −0.100, *p* = .128); There was also no significant overall difference between the control and inhibition groups, *b* = 0.075, *p* = .276. The difference in lever pressing between the two CS types did not significantly differ between groups, *b* = −0.075, *p* = .419 (Figure 4H).

#### Locomotion

STN chemogenetic inhibition resulted in greater total distance travelled, *t*(13.31) = −3.68, *p* = .003; longer total moving time, *t*(13.84) = −4.50, *p* < .001; and longer maximum movement duration, *t*(13.83) = −3.98, *p* = .001, compared with the Control group. The Inhibition group also showed a numerically higher maximum velocity (M = 115.05 cm/s) than the Control group (M = 90.57 cm/s), although this difference did not reach statistical significance, *t*(11.94) = −2.13, *p* = .055 (Figure 4I– L). Overall, STN-inhibited rats exhibited greater locomotor activity compared to control rats.

## Discussion

Previous studies have shown that STN contributes to motor, associative, and cognitive functions^25^. However, how the STN processes and integrated information to support adaptive behaviour across different learning contexts remained unclear. In this study, we combined fibre photometry and chemogenetic inhibition to investigate contribution of the STN across three behavioural paradigms: Pavlovian conditioning and its reversal, instrumental responding with its reversal and probabilistic PIT paradigm. We show that STN activity represents learned cue-reward and action-reward associations, and theses representation dynamically updated following contingency reversal. Chemogenetic inhibition revealed a contribution of the STN to behavioural flexibility and rewardguided responding. Together, our findings suggest that the STN contributes to processing and updating reward-related information, potential through integrate relevant reward associate information to guide adaptive behaviour.

### STN dynamically encodes reward and associated cues

Previous studies have identified that the STN is important for reward processing using lesion approaches^19,29^ our findings extend these finding by providing real-time evidence that the STN is integral in reward and cue recognition in established learning and is also involved in the discrimination between rewarded and unrewarded cues. Our findings demonstrate that the STN dynamically encodes both reward predictive cues and reward outcomes. During established Pavlovian conditioning, STN neurons exhibited temporally precise responses to the rewarded cue (CS+) and to reward delivery, whereas no response was observed following presentation of the unrewarded cue (CS-). Furthermore, chemogenetic inhibition of the STN impaired behavioural discrimination between rewarded and unrewarded cues. The robust STN response to reward delivery is consistent with previous fibre photometry studies reporting increased STN activity following presentation of liquid or solid foods^30^. Single-neuron recordings in non-human primate STN show reward related neuronal activity, and differences in activity between types of reward ^18,31^. Recent multielectrode recordings showed that the STN exhibited the strongest responses to reward within the basal ganglia^32^. Importantly, based on the temporal separation of neural responses to cue presentation and reward delivery shown in the present study, it seems that these events are encoded independently. This interpretation is further supported by the reversal learning response, in which STN activity continued to respond to the reward delivery with no response to the newly rewarded cue. Together, these findings establish that the STN does not simply signal reward but dynamically encodes both predictive and consummatory aspects of reward.

### STN updates reward representations and value to support cognitive flexibility

A central finding of this study is that the STN dynamically updated reward representations when cue-outcome contingencies changed. Fibre photometry recordings during reversal learning showed that STN neurons continued to respond to the previously rewarded cue (old CS+, now CS−), while failing to respond to the newly rewarded cue (new CS+). This persistence in neural activity indicates that previously learned cue-reward associations are retained within the STN and are not immediately updated following changes in reinforcement contingencies. Consistent with this interpretation, chemogenetic inhibition of the STN impaired learning of the new contingencies, demonstrating that STN activity is required for the flexible updating of cue-reward associations and adaptation to changing behavioural rules.

Previous electrode recording studies have shown that STN neurons respond to reward prediction errors and unexpected changes in reward value^33,34^, suggesting that continued activation to the former CS+ may reflect an error signal that suppresses previously reinforced responses while new cue-reward associations are acquired. Similarly, sustained STN activation has been reported during context-induced alcohol seeking and in response to previously learned drug-associated cues ^24,35^, indicating that the STN retains behaviourally relevant reward memories across diverse motivational contexts. This adaptive neural shift is also observed during the reversal of action-outcome contingencies in the instrumental conditioning paradigm where the STN neurons continued to respond to the relevant lever press on the 1^st^ day of reversal, but once the behaviour was reversed this neural response was no longer active. Together, these findings identify the STN as a dynamic substrate that continuously updates reward representations to support cognitive flexibility.

Importantly, we extend these findings across behavioural domains by showing that the STN encoded cues indicating probability of a reward. This effect has also been replicated in Parkinson’s patients, where low-frequency STN neuronal activity responded to subjective reward values and effort costs during decision making^16^. Although the animals did not show Pavlovian instrumental transfer, disruption of STN activity impaired outcome specific responses, indicating that the STN integrated reward value to guide goal-directed behaviour. This positions the STN within neural circuits underlying outcome-specific decision-making processes.

### Integration of cognitive and locomotor control

Optogenetic inhibition of STN has been shown to increase movement ^36–38^. Here, we also report hyperlocomotion induced by a chemogenetic approach. During chemogenetic inhibition of the STN during both Pavlovian conditioning and reversal, although the magazine entries during CS+ presentation decreased (supporting the role of STN in cue-reward association), the total number of magazine entries did not change. Our findings highlight a likely multi-faceted role of the STN in facilitating cognitive evaluation or other functional modalities such as motor output. This suggests a model in which learned associations and general locomotor activity are regulated by different STN mechanisms. In line with this model, single neuronal activity recording in humans and non-human primate STN show that action selection and cancellation both occur in the STN but are regulated by inputs from the globus pallidus and prefrontal motor cortex respectively^11,39,40^.

The current results suggest that the integration of cognitive and locomotor functions mediated by the STN may provide a mechanistic framework for understanding how STN deep brain stimulation (DBS) can influence both motor symptoms and cognitive-affective processes ^41–45^. Moreover, STN-DBS has been suggested as a therapeutic target for addiction ^18,46–49^. In line with this, our data provides insight into STN activity during cue-reward associations and how the STN may retain previously rewarded associated cues as a key trigger of cue or context-induced relapse to drug seeking^24,50^.

### Limitations

Several limitations should be considered when interpreting the present findings. First, although fibre photometry enabled real-time monitoring of STN population activity during reward learning, this technique measures bulk calcium dynamics and therefore cannot resolve the activity of individual neuronal subpopulations or distinguish between functionally distinct cell types. Future studies combining cell type-specific recordings or single-cell electrophysiology with behavioural paradigms will be important for determining how individual STN neuronal populations contribute to reward encoding and behavioural flexibility.

Second, although chemogenetic inhibition established a causal role for the STN in reward learning and reversal, this approach lacks the temporal precision required to dissociate the contributions of the STN during discrete behavioural events, such as cue presentation, reward delivery, or action selection. Temporally precise manipulations using optogenetic tools will be valuable for defining the specific phases of learning during which STN activity is required.

Third, despite robust Pavlovian discrimination, we did not observe Pavlovian instrumental transfer. Control animals exhibited clear conditioned responding, as evidenced by significantly greater magazine entries during the CS+ compared with the CS- and during the CS70 compared with the C30, demonstrating that the Pavlovian associations were successfully acquired. However, these conditioned cues failed to enhance instrumental responding during the transfer test. One likely explanation is that instrumental training was insufficient to establish a strong action outcome association capable of supporting transfer. Consequently, while our findings only demonstrate intact Pavlovian learning, the absence of PIT precludes conclusions regarding the contribution of the STN to the integration of Pavlovian incentive value with instrumental action selection.

Finally, viral expression was largely confined to the STN, although some spread to neighbouring structures, including the zona incerta, cannot be excluded. However, this is unlikely to explain the observed behavioural effects, as zona incerta inhibition produces outcomes opposite to STN inhibition, particularly in locomotor activity. Optogenetic inhibition of the zona incerta decreased locomotor activity, opposite to STN inhibition^49–51^. Nevertheless, greater anatomical specificity using intersectional viral strategies would further strengthen circuit-level interpretations.

## Conclusion

Collectively, these findings extend the role of the STN within the basal ganglia. Rather than functioning solely as a motor nucleus, the STN continuously encodes and updates learned reward representations that guide adaptive behaviour. These findings provide a mechanistic framework linking STN activity to reward learning, cognitive flexibility and goal-directed behaviour, with important implications for neurological and psychiatric disorders characterised by impaired behavioural adaptation.

## Acknowledgement

This work was supported by the Australian Research Council (DP220100412) to A.A.P and J.L.C. The authors declare no competing financial interests.

## Contribution

A.A.P. and J.L.C. designed the experiments; J.C and M.R. collected the data; J.C. and G.Z.Z analysed data; A.A.P, J.C, G.Z.Z and J.L.C drafted, edited and reviewed the paper.

## Discourse

We are currently consideration to upload this manuscript on an open-access online platform ( bioRxiv ) to share scientific findings rapidly for grant reviewing process.

